# Control of behavioral uncertainty by divergent frontal circuits

**DOI:** 10.1101/2024.03.29.587380

**Authors:** Yihan Wang, Qian-Quan Sun

## Abstract

Both ambiguous inference from current input and internal belief from prior input causes uncertainty. The uncertainty is typically manifested as a normal distribution at behavioral level when only current inference is manipulated as variable. When prior belief is varying, some decision relevant neural representations are dissociated. Under this circumstance, it is unclear how to describe the uncertainty and how dissociated neural representations cooperate to control the uncertainty. By simulating an unpredictable environment, which incurs conflicting valence-dependent prior beliefs, we found that a behavioral outcome, waiting time, does not follow a normal, but a log-normal distribution. By combining electrophysiological recordings, computational modeling, optogenetic manipulation, scRNA-seq and MERFISH, we showed that the formation of this behavioral outcome requires the temporally hierarchical cooperation of the neural representation of decision confidence and B230216N24Rik marked neural representation of positive and negative belief in the medial prefrontal cortex (mPFC). In summary, our results provide a mechanistic link between the dynamics of valence-dependent prior beliefs and behavioral uncertainty.

## Introduction

Behavioral uncertainty derives from the ambiguity and noise of input and prior belief of an agent. Normal distributions are frequently used to describe the behavioral uncertainty^1^. However, such distribution is conditional^2^. The normal distribution can only be applied when only one factor, such as random noise of input, is varied. In a natural environment, many external factors are unstable and unpredictable. This may cause animal’s assigned valence^3^ changes from time to time. It will further incurs instability and broader deviation of internal belief. Therefore, it is questioning about using normal distribution to describe uncertainty when both input noise and valence-dependent prior belief are varied. As the center of neural encoding of probability and uncertainty, frontal cortex circuits represent latent variables of inputs^4, 5, 6, 7, 8, 9^ and reflect internal beliefs from prior knowledge^10, 11^. When the internal belief update, different parts of frontal cortex separately reflect different information, including previous attended and newly attended information^12^. However, how the dissociated neural representations cooperate to control the behavioral uncertainty is unknown.

To test whether a behavioral outcome follows a normal distribution when faced with varying valence-dependent prior beliefs, thirsty mice were trained to lick the randomly delivered water (>75% chance) and quinine (<25% chance) in the darkness while no sensory cue indicating which liquid comes out (**Figure 1A, B; Supplemental Figure 1C**). We used waiting time from delivery onset (DO) to initiation onset (IO, the first time point when lick frequency > 6Hz) to measure behavioral uncertainty (**Figure 1B, video**). Such measurement can avoid counting occasional lick after the DO and is available in most trials (∼90%) (**Supplemental Figure 1E, F**). As a result, we found that the IO did not follow a normal distribution (P = 9.1×10^-4^) but was closer to a log-normal distribution (P = 0.4988) (**Figure 1C**).

**Figure 1.**
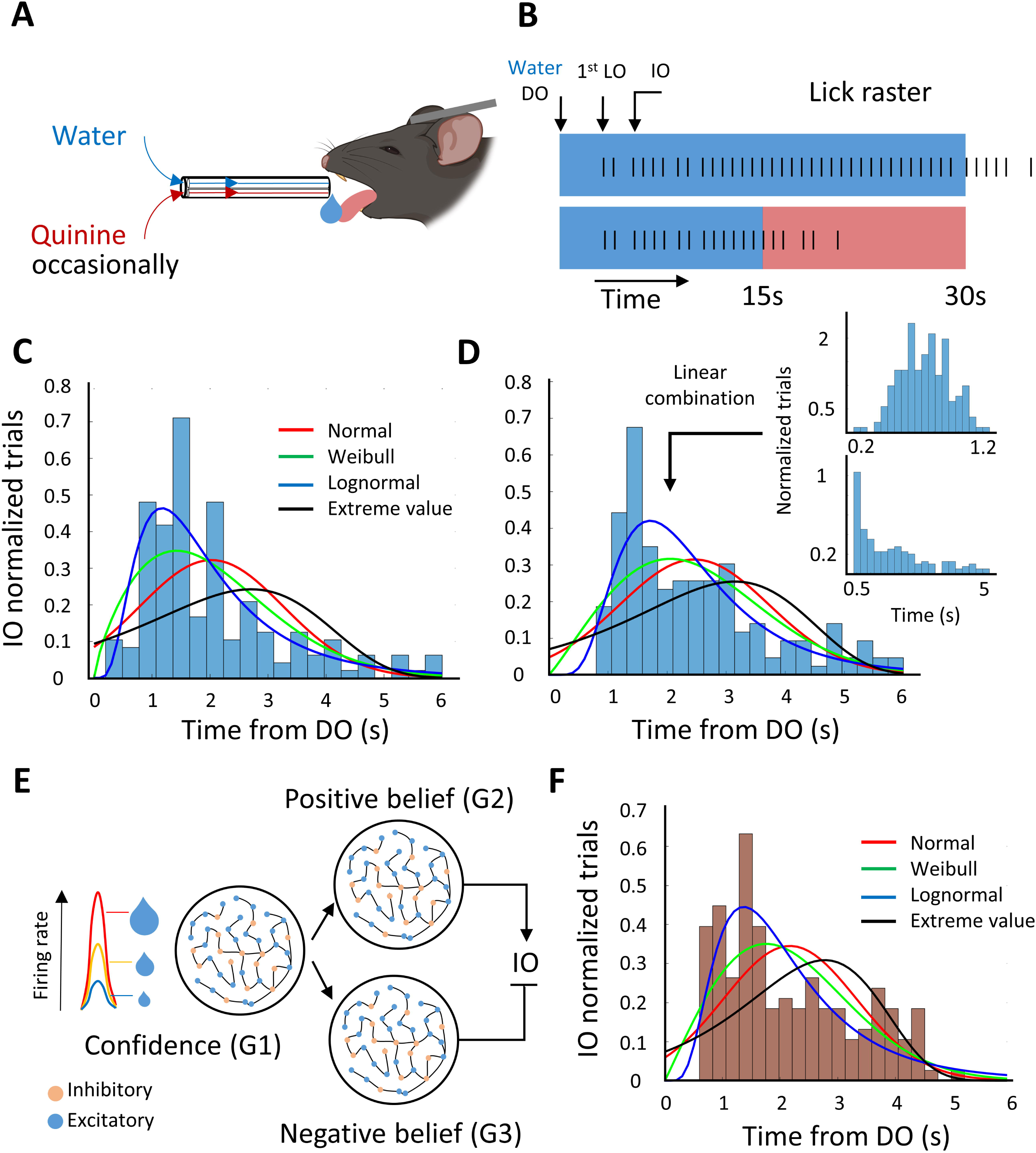
A generative model recapitulates the distribution of IO. A. Schematic images of the experimental setup (left) and the task design (right). B. Schematic lick raster plot show that only water sessions were used for recordings. Initiation onsets (first time point when the lick frequency larger than 6 Hz; IO) were measured. Abbreviation: DO: delivery onset; LO: lick onset; IO: initiation onset. C. IO distribution (165 trials, 14 mice; we did not count the trials that IO started 6s after the DO) is approximately a log-normal, but not a normal distribution. Anderson-Darling test: Normal: p=9.1×10^-4^; Weibull: p=0.0567; Log-normal: p=0.4988; Extreme value: p=3.7×10^-6^ D. The log-normal distribution was linearly combined by the normal (top right) and exponential (bottom right) distribution. Anderson-Darling test: Normal: p=0.0016; Weibull: p=0.0211; Log-normal: p=0.1717; Extreme value: p=1.7×10^-5^ E. A spike neural network model is constructed by three groups. When the firing rate of group 1 (G1) neurons reached the threshold, group 2 (G2) and group 3 (G3) neurons were activated to promote and inhibit IO, respectively. F. The IO distribution generated by the model. Anderson-Darling test: Normal: p=0.0066; Weibull: p=0.0619; Log-normal: p=0.1281; Extreme value: p=5.9939×10^-4^

In this study, we sought to understand how the different neural circuits cooperate to control this behavioral outcome. First, we created a spiking neural network model that successfully recapitulates the distribution of IO. This network consists several functionally distinct neuronal groups that cooperate in a temporally hierarchical manner. Second, we discovered two prefrontal circuits that have similar coding, functional, and collaborative properties to their counterparts in the neural network model. Thirdly, we confirmed their uniqueness from other frontal neurons by mapping their transcriptomic profiles. Finally, we demonstrated that their coding is flexible, and the log-normal distribution can be widely used.

## Results

### A generative model

Since the log-normal distribution can be a linear combination of normal and exponential distribution (**Figure 1D**), we hypothesized that there are at least two types of neuronal groups, which underlie the formation of these two distributions, respectively. The firing rate of the first group (G1) was designed to correlate with the linearly increased liquid size. Since the speed of liquid delivery across trials were given at a normal distribution, the output of this group should also follow a normal distribution. When the firing rate of the G1 reached a certain threshold (firing rate = F_0_; time = t_0_), the second group (G2) was activated to induce IO. To replicate a decreased probability at the tail of log-normal distribution, we reduced the decoding accuracy 1s after the lick onset (LO). We also added a third group (G3) to prevent the occurrence of IO immediately after the G1 reached the threshold. The G3 had the same encoding properties as the G2. In summary (**Figure 1E**; see **Supplemental Figure 2** for the details of model design), the G1 can be thought as a neural representation of decision confidence^4, 5^, which varied with reward size. G2 and G3 fulfill the following two criteria: (1) decreased decoding after t_0_+1s; (2) G2 and G3 promote and inhibit the IO, respectively. As such, G2 and G3 can be thought of as a positive and negative belief on whether to initiate a motivated behavior. Note that the positive belief is to believe that the water session will not be interrupted by quinine (top, **Figure 1B**) while negative belief is that the quinine will come (bottom, **Figure 1B**). Finally, we ran this generated network with the same trials as the experiment and acquired a similar IO distribution (**Figure 1F**). This result suggests that this triple group model captures a potential network mechanism that generates behavioral uncertainty when both decision confidence and internal belief varies.

### The role of dMP and vMP neurons in decision making

Next, we asked if the G2- and G3-like neuronal groups exist in the biological brain. In rodents, the motor cortex projecting neurons in the dorsal medial prefrontal cortex (dMP) excite motor neurons^13^ and promote the initiation of persistent lick^14^. However, silencing them does not change animals’ behavior in the middle and end phase of a persistent lick, which suggests that it’s coding may decrease after the licking initiation (criterion #1, see above generative model). The motor cortex projecting neurons in the ventral medial prefrontal cortex (vMP) also excite the motor neurons^13^ but have different projection profiles than the dMP neurons (dMP projected to dmPFC, M1, and Striatum while vMP projected to vmPFC and M1; **Supplemental Figure 3D**). In humans, the dorsal and ventral parts of the frontal cortex have been suggested to have opposite functions^15^. Above evidence suggests that vMP may have negative effect on the behavioral outcomes (criterion #2, see above generative model). Hence, we hypothesized that both dMP and vMP neurons may be the biological analogy of G2 and G3 neuronal groups, respectively. We tested this hypothesis by examining the coding of dMP and vMP neurons, and the effects of optogenetic silencing these neurons on IO.

To investigate the coding of dMP and vMP neurons, we used opto-tagging approach (**Methods**) to record the activity of dMP and vMP neurons (**Figure 2A, B; Supplemental Figure 4A, B**) while thirsty start to lick the water. Our results showed that both dMP and vMP neurons responded exclusively (significantly increased mean neuronal activity after the LO, **Figure 2F**) and repeatedly (significantly higher decoding accuracy than the pseudo data) after the first lick onset (LO) (P1, **Figure 2J**). Although their firing rates remained at a high level in all recorded periods as population (blue traces, **Figure 2F**), the firing rate of individual dMP and vMP neurons became more various across trials after LO+1s than P1 (LO to LO+1s) (**Figure 2D, E**). This was indicated by the lower decoding accuracy (**Figure 2J**). Meanwhile, the G2 and G3 in the model captured similar results to dMP and vMP neurons (**Figure 2G, H, I, and K**). These results suggest that two types of MP neurons reflect the coding properties of the G2 and G3 in the model (criterion #1).

**Figure 2.**
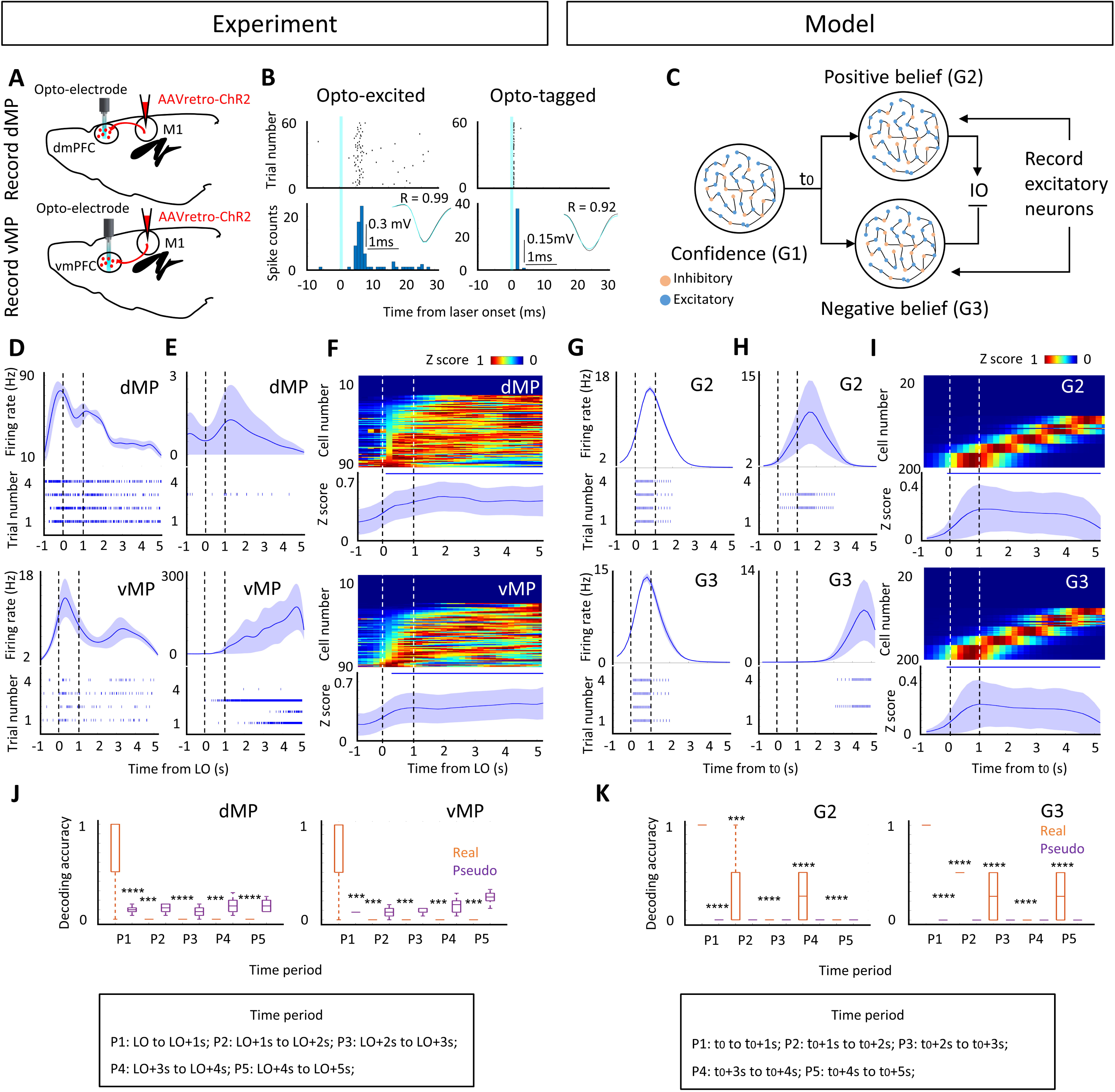
Two MP neuronal subtypes have similar coding properties with the model. A. Schematic images showing the recording of dMP::ChR2 (top) and vMP ::ChR2 (bottom) neurons. B. Identification of MP neurons. Both opto-excited and opto-tagged units were identified as MP neurons. C. Schematic images showing the excitatory neurons from G2 and G3 were recorded. D & G. Firing of representative dMP (top, D), vMP (bottom, D), G2 (top, G), and G3 (bottom, G) neurons showing persistent and excitatory activity from trial to trial in the P1 (LO/t0 to LO/t0+1s). E & H. Firing of representative dMP (top, E), vMP (bottom, E), G2 (top, H), and G3 (bottom, H) neurons showing various firing from trial to trial in the P2 to P5 (LO/t0+1s to LO/t0+5s). F & I. Color coded maps showing normalized activity of individual dMP (top, F), vMP (bottom, F), G2 (top, I), and G3 (bottom, I) neurons. Blue traces showing the mean activity of these neurons. Shade represents ±S.E.M across the trials. The horizontal bar represents the significantly (p<0.05) higher activity than the baseline (LO/t0-1s to LO/t0). J & K. Decoding of the time epochs after the 1^st^ LO or t0 using the normalized activity of dMP (left J), vMP (right J), G2 (right K), and G3 (right K) neurons. Two sample t-test was used to compare P1 pseudo, P2, P3, P4, and P5 with P1 data. Asteroid represents significantly lower decoding accuracy than P1 data. ***p<0.001, ****p<0.0001.

To study the functional similarity between MP neurons and the model, we examined whether dMP and vMP neurons promote or suppress the initiation of licking actions. We expressed stGtACR2 in the MP neurons and shined laser to manipulate them (**Supplement Figure 4C, D**). As expected, opto-silencing of dMP neurons delayed IO (ibias, equivalent to 1/IO: no laser > laser; **Figure 3A, C**), whereas inactivation of vMP neurons increased action initiation (ibias: no laser < laser; **Figure 3B, D**). However, inactivation of either dMP or vMP neurons did not affect (dMP: p=0.4831; vMP: p=0.6863) behavioral extinction triggered by quinine stimuli (**Supplemental Figure 5**), suggesting that the initiation and termination of a persistent licking action are controlled by different brain circuits. In addition, different from dMP neurons, parts of the vMP neurons (1% dMP vs 18% vMP) responded to quinine stimuli during the action extinction period (**Supplemental Figure 6**). It suggests that a negative belief about initiating a persistent action is encoded and stored in the vMP neurons. Collectively, our results indicate that dMP neurons promote the initiation of persistent lick, whereas vMP neurons have a negative effect on it (criterion #2).

**Figure 3.**
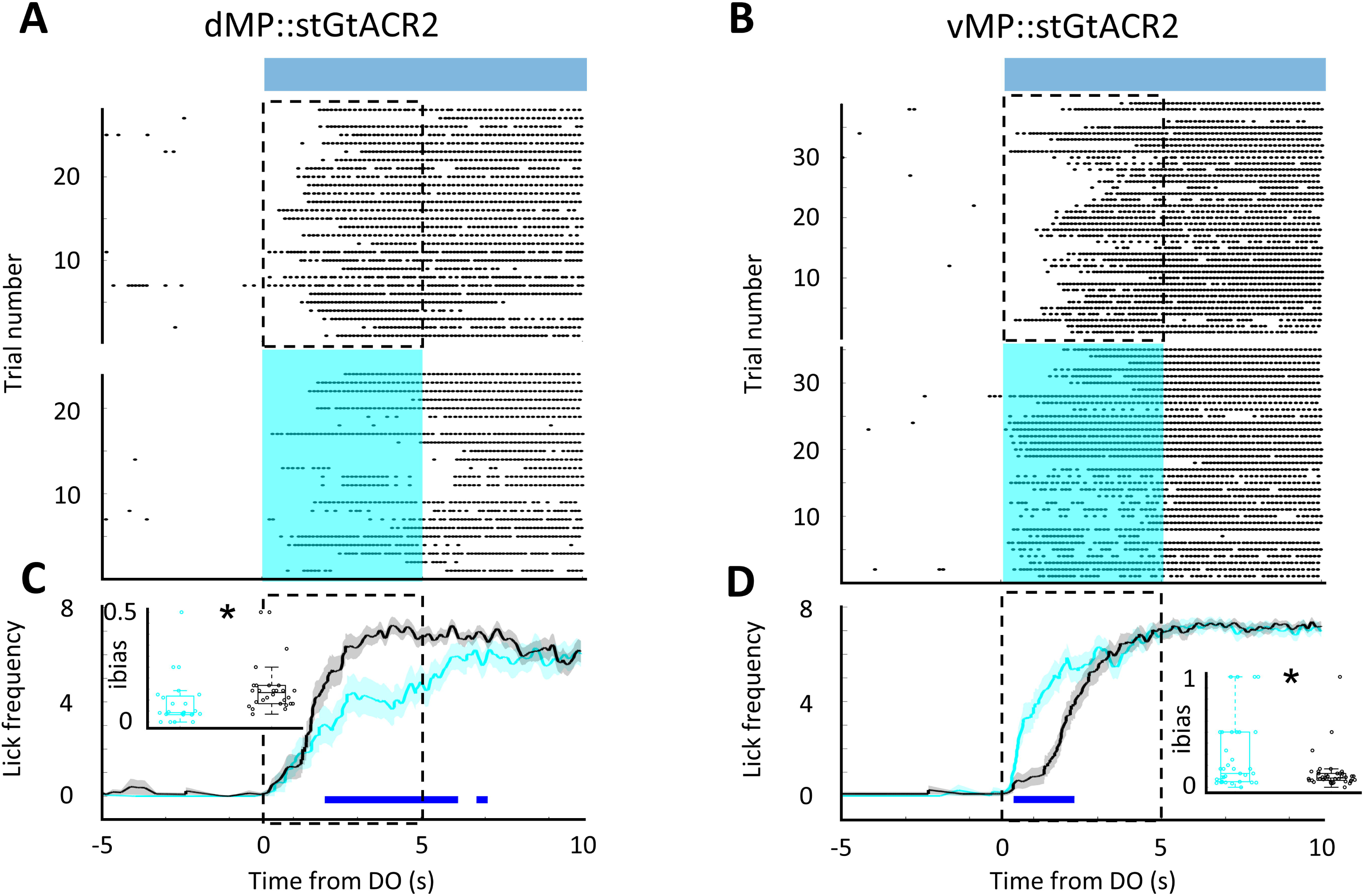
Two MP neuronal subtypes show opposing function toward initiation of motivated behavior. A-B. Licking activity from dMP::stGtACR2 (A) mice (n=4) and vMP::stGtACR2 (B) mice (n=4) without laser (top) and with laser (bottom). Blue bar represents the period of water delivery. C-D. The traces of lick frequency and the box plot of initiation bias (ibias, equivalent to 1/IO, inset) showing that inactivation of dMP (C) and vMP (D) neurons significantly decrease and increase the initiation of persistent lick, respectively. The trace shade represents ±S.E.M across the trials. Horizontal bar represents the significantly different lick frequency (p<0.05). Laser: cyan; no laser: black. Statistics: two sample t-test, *p<0.05.

Therefore, the above results suggest that dMP and vMP neurons constitute the G2- and G3-like neuronal groups described in our generative model. Demonstrated by the neural network model and the empirical data, dMP and vMP neurons contribute to the formation of a log-normal distribution of IO.

### Transcriptomic profiles of mPFC MP neurons

In the mPFC, MP neurons exhibit heterogeneously electrophysiological properties and distribute unevenly across layers^13^. To further investigate the unique transcriptomic properties of MP neurons, single-cell RNA sequencing (scRNA-seq) and multiplexed error robust fluorescence in situ hybridization^16^ (MERFISH) were used to identify their transcriptomic profiles and marker genes. First, we labeled MP neurons by injecting retro-AAV into the motor cortex. Then, all cells were extracted from the mPFC and their transcriptome was examined using scRNA-seq, and Allen data^17^ registering (**Supplemental Figure 7A**). Consistent with our previous findings that MP neurons project to cortex and striatum instead of deep brain regions^13^, sequencing data showed that majority (88%) of MP glutamatergic neurons were classified into intratelencephalic (IT), instead of pyramidal tract (PT), cluster (**column 2 Figure 4C**). Surprisingly, the majority (70%) of Bcl11b-positive (preferred transcription in PT neuron^18, 19^) MP glutamatergic neurons were also classified into IT cluster (**column 2 Figure 4C**). Next, random sample clustering (**Methods**), MERFISH, and single-cell graph cuts optimization^20^ (scGCO) were used to filter the marker genes of the MP neurons (**Supplemental Figure 7B, C and 8A**). Specifically, the candidate marker genes were first extracted by random sample clustering. Then, scGCO was used to remove the marker genes without spatial bias on MERFISH slices. Finally, the selected genes were further compared their spatial distribution with MP neurons. As a result, B230216N24Rik gene was selected with the most similar spatial distribution with MP neurons among all MP neuron marker genes (**Figure 4A, B; Supplemental Figure 8B, C**). The similarity of transcriptomic profiles between B230216N24Rik-positive and MP glutamatergic neurons was also confirmed by scRNA-seq data (**Figure 4D, E; Supplemental Figure 8D**). These results indicate that MP neurons are transcriptomic unique and marked by B230216N24Rik in the mPFC. It, thus, may genetically distinguish MP neuron from the neural representations of decision confidence in the mPFC. However, B230216N24Rik and other marker genes of MP neurons did not show the apparent discrimination of dMP and vMP neurons (**Supplemental Figure 9**). It suggests that the transcriptomic distinction between dMP and vMP neurons may be described by specific spatial marker genes in the mPFC.

**Figure 4.**
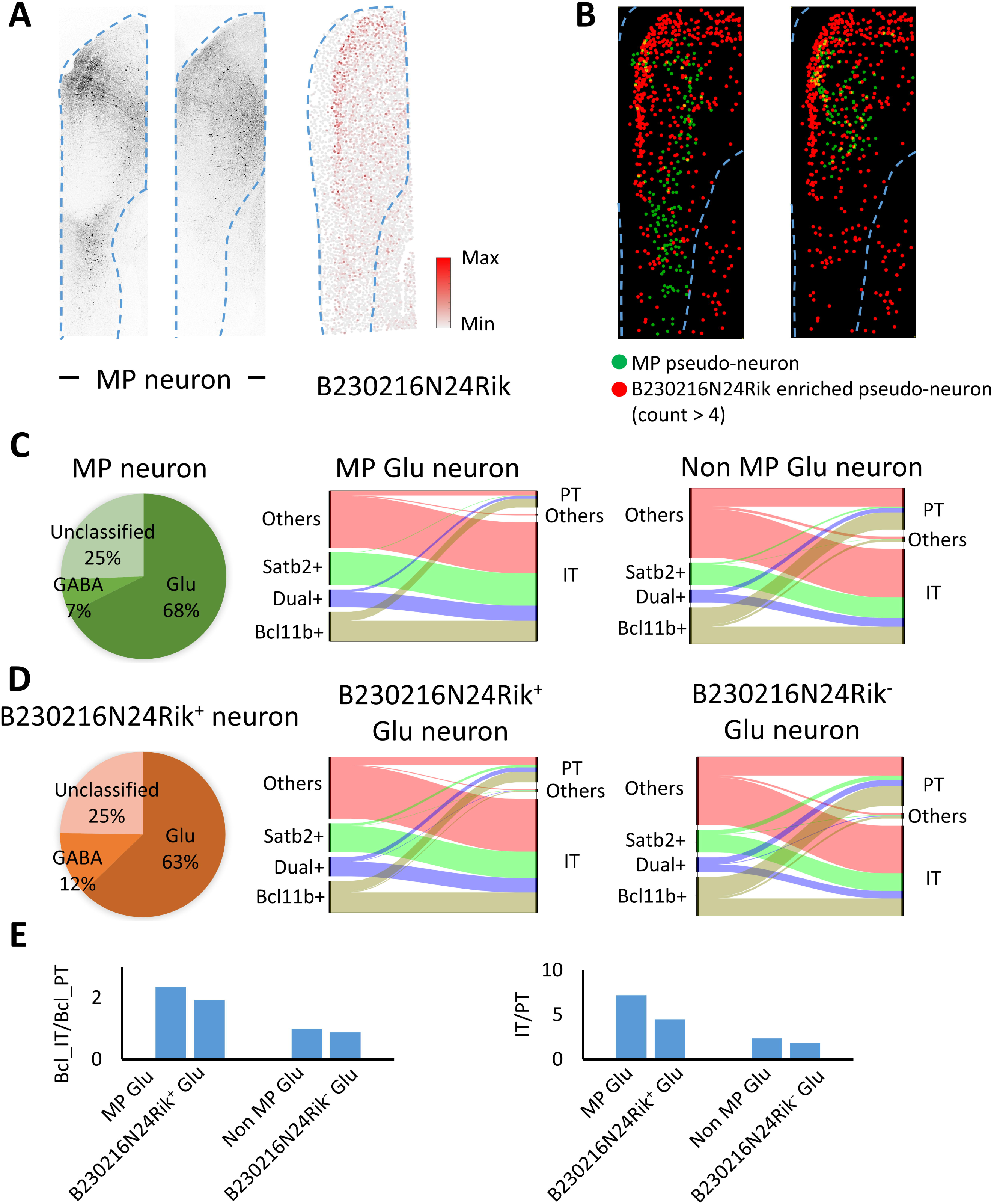
MP neuron in the mPFC can be marked by B230216N24Rik. A. The distribution of MP neuron and B230216N24Rik transcripts is shown in the Zeiss and MERFISH slices, respectively. B. The MP neuron and B230216N24Rik enriched neuron have similar distribution in the mPFC. To visualize those two types of neurons in a standard way, the Zeiss slices with MP neuron were registered into the MERFISH slices with B230216N24Rik transcriptome and then all neurons were labeled by equal sized circles (pseudo-neuron). C-E. MP neuron and B230216N24Rik positive neuron have similar transcriptomic profile which revealed by scRNA-seq. The transcriptomic profile of MP neuron (C) and B230216N24Rik positive neuron (D) is shown in the broad neuronal type (left pie chart) and glutamatergic type (right alluvial plots). Their comparisons are shown in the bar graph (E).

### Amplified MP neuronal activity across training

In the prefrontal cortex, projection neurons amplify or decrease their activity across the training in response to the conditioned stimulus^21^. Since dMP and vMP neurons are projection neurons in the prefrontal cortex, we hypothesized that they also follow this principle. To examine the plastic profile of these two types of neurons, we recorded their activity before and after training in the task (**Figure 5A**). In vivo electrophysiological recording revealed that the activity of both dMP and vMP neurons were significantly amplified (dMP p<0.01; vMP p<0.0001) specifically in response to the water delivery (1-3s after water delivery onset) across training (**Figure 5B, C**).

**Figure 5.**
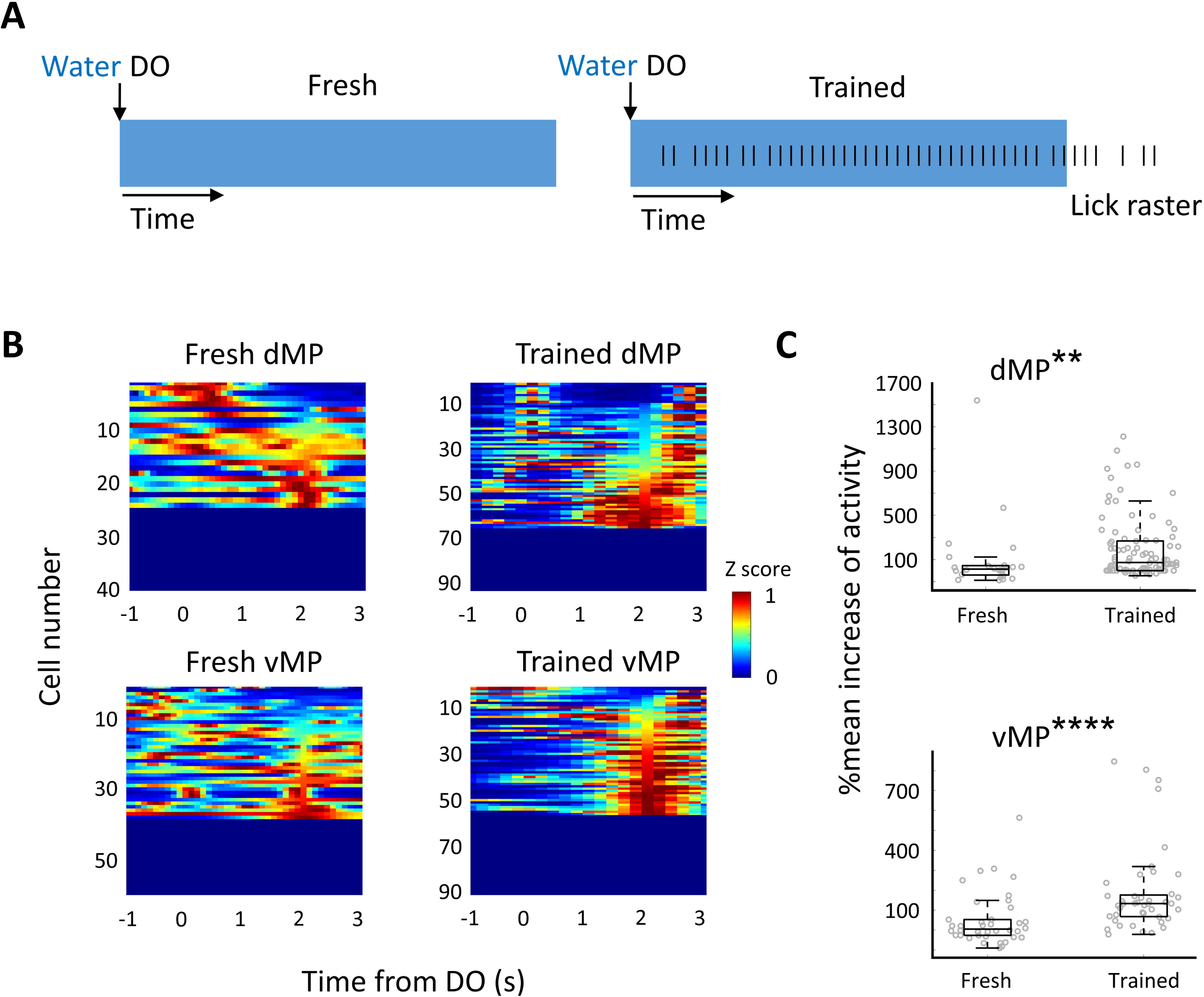
Response of two MP neuronal subtypes to the water delivery are amplified across training. A. Schematic images showing licking response to the water delivery in the fresh (left) and trained (right) trials. Fresh: no lick during the water delivery period; Trained: the persistent lick time period, of which the lick frequency is more than 6Hz, exceeded 2/3 of water delivery time. B. Color-coded map showing normalized activity of individual dMP (top) and vMP (bottom) neurons in the fresh (left) and trained (right) trials. C. Box plot showing that dMP (top) and vMP (bottom) significantly increased their activity across training. Mean increase of activity was calculated as the averaged firing rate (from DO+1s to DO+3s) increased from the baseline (from DO-1s to DO+1s). Statistics: Kolmogorov-Smirnov test **p<0.01, ****p<0.0001.

Next, we sought to understand if such neural activity amplification happens in other situations that lack opposing stimulations. The mice were repeatedly administered sucrose or shock. The neuronal activity of the MP neurons and their marker gene enriched neurons in the first trial (fresh emotional experience) and in the 5^th^ or 6^th^ trial (trained) were measured. Enabled by scRNA-seq and neural activity labeling^22^ on TRAP2-Ai9 mouse line (**Supplemental Figure 10A**), we extracted the marker genes of activated neurons in response to sucrose or shock stimuli (**Supplemental Figure 10B-D**). Then, we extracted MP neurons, MP neuronal marker gene enriched neurons, TRAP2 neurons, and TRAP2 neuronal marker gene enriched neurons. We found that MP neurons and their marker genes enriched neurons were not activated in fresh emotional experience (i.e., mixed of foot shock and sucrose activated neurons and their marker gene enriched neurons did not overlap with MP neurons and their marker genes enriched neurons; **Supplemental Figure 11A, B**). However, both dMP and vMP neurons increased their activity after 5 to 6 times training with sucrose or foot shock stimuli (i.e., more dMP and vMP neurons merged with activated neurons; **Supplemental Figure 11E, F**). These results indicate that both dMP and vMP neurons amplify their activity across training in response to either positive or negative emotional stimuli (i.e. both dMP and vMP neurons do not encode valence). We suspected that such response was due to the influence of other environmental factors on internal beliefs that are encoded by dMP and vMP neurons. Moreover, the fitness of log-normal distribution was always better than that of normal distribution when the weights of dMP and vMP counterparts (G2 and G3) were adjusted in the model (**Supplemental Figure 12**). Collectively, our results suggest that the log-normal distribution of IO can be applied to more general situation where the ratio of conflicting prior stimuli can be ignored.

## Discussion

In summary, we demonstrated that the behavioral outcome cannot be simply described by normal distribution when both decision confidence and internal belief varies. In the frontal cortex, it requires a hieratical cooperation among multiple heterogeneous neuronal groups to fully account for the behavioral uncertainty. Therefore, behavioral outcomes can be more flexible rather than simply following a normal distribution.

In detail, we explained in an example of how multiple neuronal groups hieratically collaborate with each other to perform a task. First, the neural representation of decision confidence is activated to respond external stimulation. When its firing rate reaches a threshold, a certain event (this could be the first lick after DO or a facial movement in our case) is induced. This event further excites downstream dMP and vMP neurons in the mPFC. They modulate the initiation of a motivated behavior positively and negatively. The coding of dMP and vMP neurons decreases along with the time proceeding. As a result, the waiting time to accept the reward is log-normally distributed.

In consistent with our previous finding^14^, we found that both dMP and vMP neurons do not represent valence. Instead, they encode the positive and negative belief about whether to initiate motivated behaviors. As the activity of both dMP and vMP neurons amplify across either repeated sucrose or shock stimuli, such belief can happen under either positive or negative valence. For example, during water delivery, both positive and negative beliefs about whether to accept it or not will be generated in the animal’s brain.

High degree of uncertainty about future event may lead to anxietie^23, 24^. Additionally, growing evidence reported the deficit in prefrontal cortex are linked to elevated anxietie^25^. Since the activities of dMP and vMP neurons contribute to behavioral uncertainty, the transcriptomic profile of MP neurons may provide a potential target for treatment of condition associated anxieties. Recovering the function of subtypes of MP neurons may reduce anxiety. However, further work is needed to analyze the genetic profiles specific to dMP or vMP neurons.

## Methods

### Subject details

All experimental procedures were approved by the Institutional Animal Care and Use Committee (IACUC) and the University of Wyoming Institutional Biosafety Committee (IBC). All mice, including WT, Ai9, and Ai9-Trap2, were bred on C57/BL6J background. Immunocompetent mice were used for various experimental purposes. Mice were housed alone in a vivarium at 21-23 °C and a 12-h light/dark cycle at least 7 days before implantation or water deprivation. For electrophysiological and optogenetic silencing experiments, 9 female and 5 male mice were used. For single cell sequencing, 20 female and 11 male mice were used. 4 female and 2 male mice were used in preparation of Multiplexed Error-Robust Fluorescence in situ Hybridization (MERFISH) slices. 6 female and 5 male mice were used for Zeiss imaging to measure axon length and count cell number. We did not find significant difference between female and male mice on the behavioral task.

### Surgeries

For all surgeries, mice were anesthetized with 2% isoflurane (2 LPM for induction and 0.4 LPM for maintenance). Their heads were fixed to the stereotactic device (NARISHIGE SG -4N) and their body temperature was maintained with a heating pad (K&H No. 1060). Seventy percent isopropyl alcohol followed by iodine was applied to the incision site. To expose the skull, the skin was incised, and the dura was removed. The coordinates used for positioning the injection and implantation sites were relative to Bregma (antero-posterior A-P, medio-lateral M-L, dorsal-ventral D-V) in mm. We used sterile surgical techniques (all surgical instruments were sterilized before the operation) and injected intraperitoneally 50 mg kg^-1^ ibuprofen for postoperative recovery.

P14-30 mice were used for viral injection. A small craniotomy (approximately 0.2 mm in diameter) was made over the injection site. Glass filaments (Drummond Scientific Co.) with a tip diameter of approximately 5 µm were filled with 2 µL of virus solution. By pressure injection with a custom-made device driven by a single-axis hydraulic manipulator (NARISHIGE mmo-220A), the virus solution (undiluted, 100 nL at each injection site) was delivered to the desired regions at a rate of 30-50 nL min^-1^. To label MP neurons in the mPFC, pAAV-CAG-EGFP (Addgene_37825-AAVrg) was injected into the bilateral motor cortex (A-P -0.6, M-L ±1.0, D-V 0.2 0.5 0.8). To optogenetically label MP neurons in the mPFC, pAAV-CAG -hChR2-mcherry (Addgene_28017-AAVrg) was injected into the unilateral motor cortex (A-P -0.6, M-L 1.0, D-V 0.2 0.5 0.8). To optogenetically inactivate MP neurons in the mPFC, pAAV-CKIIa-stGtACR2-FusionRed (Addgene_105669-AAVrg) was injected into the bilateral motor cortex (A-P - 0.6, M-L ±1.0, D-V 0.2 0.5 0.8). Mice were returned to the home cage at least three weeks before implantation.

Aged 2-3 months mice were used for implantation. For both opto-electrode and optic fiber implantation, we made a craniotomy (the size was based on the specific opto-electrode and optic fiber) over the implant site. A customized head bar (github.com/ywang2822/Multi_Lick_ports_behavioral_setup) was placed over the skull using Metabond (C&B Metabond, Parkell) and dental cement (Lang Dental). We used silicone sealant (kwik-cast, world precision instrument) to cover exposed brain tissue. To implant opto-electrode, an optic fiber (MFC_200/245-0.37_2.0mm_MF1.25_FLT) were fixed around 0.5 mm above a silicon probe (A4×8-Edge-2mm-100-200-177-CM32, NeuroNexus) using crazy glue and dental cement (Lang Dental). This custom-made opto-electrode was then implanted into unilateral dmPFC (A-P 1.0-1.7, M-L 1, D-V 1) or vmPFC (A-P 1.0-1.7, M-L 0.5, D-V 2.5). An anchor screw was placed on the contralateral cerebellum to connect ground wires of the electrodes. To implant optic fibers, two optic fibers (MFC_200/245-0.37_2.0mm_MF1.25_FLT, Doric Lenses) were placed on the bilateral vmPFC (A-P 1.0-1.7, M-L ± 0.5, D-V 2).

### Behavioral task

The behavioral set-up and recording signals were described previously^14^. In brief, mice were head-fixed and allowed to lick different liquids, which delivered by a multi-lick-ports (github.com/ywang2822/Multi_Lick_ports_behavioral_setup), in a 30-second period. The delivery speed was manually calibrated to a certain value between 0.15 - 0.2mL min^-1^ (∼N(0.175,0.025)) every time before the behavioral test. In most cases, 30-second water was delivered. In some cases, 15-second water was delivered followed by 15-second quinine. The liquid was gradually and non-stop delivered at a fixed speed. In this case, the mice confidence to the liquid delivery increased with the time evolving. Zero time interval was set when switched to a different type of liquid. On this set-up, mice were able to freely move their limbs. Frames, speed, lick onsets (LOs), laser delivery onsets, liquid delivery onsets (DOs), and neural activity were recorded simultaneously, and the signals were sent to an USB interface board (Intan Technologies, RHD).

Before performing the task, mice were deprived of water until their body weight decreased by approximately 22%. Specifically, we first examined the duration of water deprivation for each mouse. After body weight decreased by 22%, the mice had free access to water for at least five days. We observed the mice’s health continuously. After following this procedure two to three times, the mice became accustomed to water deprivation and no obvious suffering was observed. Then, the tasks were performed.

### Initiation/termination bias and persistent period of the behavior

We calculated the initiation and termination bias (i/tbias^14^) to assess whether the mice were starting or stopping the continuous licking behavior. For both initiation and termination bias, simple moving averages (SMAs) were calculated in a 1-second time window as follows:

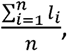

where *l* is the lick times during a 200ms time window and *n* = 5 . We created a vector that contained SMAs sampled at 200ms intervals. i/tbias were calculated as 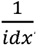, where idx is the index of the SMAs at a given time point. For ibias, the given time point was the first time that SMA > 1.2 (6Hz). This given time point was equivalent to the initiation onset (IO; IO=idx/5). SMAs were calculated from the onset of water delivery. If IO was larger than 6 (6s after DO), ibias was calculated as zero. For tbias, the given time point was the first time that SMA < 1 (5Hz). This given time point was equivalent to the termination onset (TO; TO=idx/5). SMAs were calculated from the onset of quinine delivery or the end of water delivery (water DE). The persistent licking period from water DO to water DE was calculated by adding up all the persistent inter-lick intervals, which are shorter than 200ms.

### Network modelling

BRIAN (brian2.readthedocs.io) was used to simulate the network. The network model consisted of three groups (G1, G2, and G3). In all three groups, there were 200 excitatory neurons and 50 inhibitory neurons. The neuronal equation and the synapse equation were set based on a previous work^27^. For G1, the input was given as:

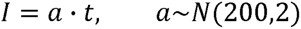

10 excitatory neurons were randomly selected. When their average firing rate reached 10 Hz, the corresponding time point was used for activating G2 and G3. For G2 or G3, we divided it into 5 subgroups, which resulted in a total of 33 neurons in each subgroup. In the first subgroup, 20 neurons were selected. In the rest subgroups, 7-14 neurons were selected in each group. We gave 76.5 to 102 pA input to the selected neurons in G2 and 75 to 100 pA input to the selected neurons in G3. Their input patterns were shown in **Supplemental Figure 2**. Then we selected all neurons in the first subgroups and randomly selected 7-14 neurons in each of the rest subgroups. Averaged firing rates of these selected neurons were calculated. The IO+0.5 was computed as the first time point when the average firing rate of selected neurons in G2 – the average firing rate of selected neurons in G3 larger than a threshold (0.9). In other simulations (**Supplemental Figure 12**), we assigned G2 and G3 to different weights by adjusting this threshold from 0.5 to 1.7.

### Single-unit extracellular recording and firing rate estimation

Signals from opto-electrodes were acquired using 32-channel RHD head stage, which connected with a signal acquisition system (USB board, Intan Technologies). We processed signals using an offline software Spikesorter^28, 29, 30^ (swindale.ecc.ubc.ca/home-page/software/). For signal filtering, 0.5 kHz and 4 poles high pass Butterworth filter were used. For event detection, median noise calculation and 80µV, 5x noise, 0.75ms window width threshold were used. For clustering, 2 PCA dimensions, -0.8 to 0.8 template window, 5 starting sigma, and 9 thresholds were used. Other parameters followed the default settings of the software. After single-unit data was extracted, we used Bayesian adaptive kernel smoother^31^ with 100 or 200ms bandwidth to estimate firing rate.

### Optogenetic silencing and opto-tagging

To optogenetically silencing stGtACR2 labeled vMP neurons, we illuminated bilateral ventral prefrontal cortexes with a 473-nm 5mW laser at a frequency of 20 Hz and a width of 40ms for 5 seconds. The onset of laser pulses was triggered based on water DO, water DE, or quinine DO. The optogenetic silencing experiments were only performed on the well-trained mice (persistent licking period from water DO to water DE > 0.66*total time period from water DO to water DE). To identify ChR2 labeled vMP and dMP neurons, we applied 473-nm 7mW laser pulses (1ms width at 20 Hz, 3s duration) on the unilaterally prefrontal cortex of viral (AAV-ChR2) injected mice. An opto-tagging approach was used as previously described^32^. In brief, laser evoked spikes and network excited spikes were assessed in a 0-6ms and a 7-50ms temporal windows after the laser onset, respectively. For those units with significant correlation (correlation coefficient > 0.80) of average waveform and significantly different distribution (P < 0.05) of spike latency with baseline units were mostly identified as laser evoked or network excited units. We also manually reviewed classified units.

### Cell classification

We classified firing rate data to discriminate initial and terminal phase of the licing movement according to our previous work^14^ and i/t bias. We used 1^st^ licking onset (LO) as the starting point for the initial phase and termination onset (TO), i.e., the first time point when the licking frequency blow 5Hz, as the starting point for the terminal phase in both the water-water and water-quinine sessions. The firing rate data at the 2-second temporal window (5 data points) after the 1^st^ water LO or TO was used. The pseudo-data was created from randomly shuffled data in the 70-second temporal window. The initial phase tuned neuron or the neural representation of initial phase was defined when it’s firing rate higher than 65% pseudo-data at the 1^st^ water LO: 1^st^ water LO+2s temporal window and lower than 35% pseudo-data at the TO: TO+2s temporal window; the terminal phase tuned neuron or the neural representation of terminal phase was defined when it’s firing rate higher than 65% pseudo-data at the TO: TO+2s temporal window and lower than 35% pseudo-data at the 1^st^ water LO: 1^st^ water LO+2s temporal window.

### Population decoding

We used a multiclass linear discriminant analysis (LDA), which was based on an error-correcting output codes (ECOC) classifier, to decode movement phases (initial and terminal phase). Similar with **Cell classification**, the zero time point of the evaluation window was aligned to the 1^st^ water LO/TO. To normalize firing rate data, we first created a shuffled firing rate matrix by randomly extracting firing rate data from whole 70s long spike train. Then the real firing rate data were normalized by dividing mean shuffled firing rate. To create firing rate pseudo data, we randomly selected zero time point from whole 70s long spike train. This procedure was repeated 50 times. Before training decoder, we reduced the dimensionality of the firing rate data by PCA to improve decoding performance^33^. The dimensions that explained over 85% of the data variance were selected to train a decoder. We used 50 percent of real and pseudo data to train the decoder, and 50 percent of data were used to test the decoder’s performance. This procedure was repeated 10 times to get an average accuracy and standard deviation.

To test whether the neural response is repeatable after the 1^st^ water LO, we trained the LDA decoder to predict time epochs using normalized firing rates. The procedure for normalizing firing rates and creating pseudo data for firing rates is the same as for decoding movement phases. The normalized firing rates were further z-scored by dividing the maximum firing rate value of all the time epochs tested. Fifty percent of the real and pseudo data were used to train the decoder, and the remainder was used to test prediction accuracy. We repeated this procedure 10 times to determine the average accuracy and standard deviation.

### TRAP2 induction and labeling activated neurons

We took advantage of TRAP2^22^ that enables highly efficient labelling of activated neurons to specifically detect the neuronal activity in response to hedonic and aversive stimuli. To visualize activated neurons, we first intraperitoneally administrated 50 mg Kg^-1^ 4-hydroxytamoxifen (4-OHT; Sigma, Cat# H6278) into TRAP2/Ai9 mouse line. 4-OHT was prepared according to a previous work^34^. Briefly, 10 mg 4-OHT was dissolved in 0.5 mL ethanol at 50 °C to acquire a concentration of 20 mg mL^-1^ and was then aliquot (100 µL/tube) and stored at -20 °C. Before use, 100 µL 4-OHT was mixed with 300 µL oil mix (a 1:4 mixture of castor oil: sesame oil) and the ethanol was evaporated by centrifugation. For hedonic stimuli, TRAP2/Ai9 mice with overnight water deprivation were fed with 5% sucrose. For aversive stimuli, two to three times 1s, 20 volts electrical shocks were delivered by a customized shocker (electric shock box machine kit, STEREN). Mice without screaming or running during the shock were excluded. For both of stimuli, aged 1-3.5 months mice were placed into a dark box with a metal grid bottom and were stimulated 15 minutes after the administration. After stimuli, mice were kept in the dark box for at least 24 hours to allow tamoxifen fully metabolized. After at least seven days, the mice were sacrificed for single-cell sequencing, or in situ hybridization, or imaging.

### Histology for Zeiss imaging

P30-P90 viral injected mice were anesthetized with 2% isoflurane (v/v) and perfused intracardially with 0.9% saline followed by 4% paraformaldehyde (PFA, pH = 7.4). Brains were extracted and stored in 4% PFA at 4 °C. Fixed brain tissues were washed three times using phosphate buffered saline (PBS) and then were dehydrated in 30% sucrose for 24 hrs. Dehydrated brain tissues were embedded with optimal cutting temperature compound (O.C.T, Tissue tek) and then were sectioned into 70 μm thickness slices. Fluorescent images were acquired by an LSM 980 microscope (Zeiss), with a 10 × 0.45NA objective or a 2.5 × 0.085NA objective.

To count the number of cells in the Zeiss images, we used a cell segmentation method. First, the edges were cleaned, and noise was removed. Holes in objects were filled and connections between objects were removed. Then, a threshold was set for the size of the objects. The objects that were larger or smaller than this threshold was selected as cells. We used different thresholds for AAV-retro and DAPI-labelled cells. Another manual correction was performed to confirm the cell count. We manually counted the cells in the images where the objects could not be detected using the ImageJ plugin Cell Counter.

To measure the axon length, we first divided the image into small parts and used our previously trained classifier^13^ to extract axons. The images with extracted axons were further registered to the Allen Mouse Brain Common Coordinate Framework v3 (CCFv3) using manually selected reference points (**Supplemental Figure 1**). The depth of dmPFC was set as 2mm from the pia and the range of vmPFC was from 2mm to 3.1mm.

### 10x Single-cell Sequencing

The procedure of sample collection was adapted from previous works^13, 17^. Briefly, aged 2-3 months mice were anesthetized with 2% isoflurane (v/v) and intracardially perfused with a cutting buffer (2.5 KCl, 1.25 NaH_2_PO_4_, 10.0 MgCl_2_, 0.5 CaCl_2_, 26.0 NaHCO_3_, 11.0 glucose, and 234.0 sucrose in mM; bubbled with 95% O_2_ and 5% CO_2_). Coronal sections were cut in 400 µm thickness by a vibrating blade microtome (Leica VT1000s). Then, these sections were transferred to 34 °C oxygenated (95% O2 and 5% CO2) ACSF (126.0 NaCl, 2.5 KCl, 1.25 NaH_2_PO_4_, 1.0 MgCl_2_, 2.0 CaCl_2_, 26.0 NaHCO_3_ and 10.0 glucose in mM) for 1 hr. mPFC was then micro-dissected from the sections. The profiling and boundaries of mPFC were defined CCFv3.

We isolated single cells from the dissected mPFC sections using a neural tissue dissociation kit (Cat# 130-092-628, Miltenyi Biotec). After cell concentration quantified, samples were loaded onto 10x genomics Chromium controller or Chromium X. For the cell capture, barcoding, reverse transcription, cDNA amplification, and library construction, we followed the instruction of Next GEM Single Cell 3’ reagent kits v3.1 Dual Index (10x genomics).

The constructed 10x libraries were sequenced on Illumina NextSeq1000 (28 cycles Read 1; 10 cycles Read 2; 10 cycles Read 3; 90 cycles Read 4). The sequencing reads were aligned to a customized mouse reference transcriptome (GRCm39 with EGFP and tdTomato mRNA sequence) using 10x Cloud Analysis. In each 10x run, cells that met the following criteria were filtered out: cells with fewer than 1500 detected genes, non-neuronal cells (RNA_Map2 < 1), and striatal cells (with mRNAs Drd2, Adora2a, and Six3). Doublets were identified and removed using DoubletFinder (https://github.com/chris-mcginnis-ucsf/DoubletFinder). Predicted gene models (gene name starts with ‘Gm’), mitochondrial chromosome genes (gene name starts with ‘Mt’ or ‘mt’), ribosomal genes (gene name starts with ‘Rps’ or ‘Rpl’), and sex-specific genes (gene name starts with ‘Usp9 or Eif2s3 or Uty or Dbx’) were removed. Cells were then classified into broad class of excitatory (RNA_Slc17a7 ≥ 2 & RNA_Slc6a1 ≤ 1), inhibitory (RNA_Slc17a7 ≤ 2 & RNA_Slc6a1 ≥ 1), and unclassified neurons.

### Random sample clustering

Since the number of EGFP and tdTomato labeled cells were far fewer than unlabeled cells, directly clustering among them are unable to extract marker genes. To solve this problem, we randomly extracted equal number of unlabeled cells with labeled cells (bootstrap) and performed an automatic iterative clustering^17^ using scrattch.hicat (https://github.com/AllenInstitute/scrattch.hicat). The parameters were set as: q1.th = 0.4, q.diff.th = 0.7, de.score.th = 150, min.cells = 4. This procedure was repeated 100 times to generate 100 marker genes files. Marker genes that were selected over 15 times from all pairwise clustering was identified as the Differential Expression (DE) genes of EGFP or tdTomato labeled cells. These DE genes were selected for subsequent multiplexed error-robust fluorescence in situ hybridization^16^ (MERFISH).

### Comparing with Allen data

To classify the EGFP and tdTomato labeled cells into layer and projection specific neuronal group, we compared the sequencing data with Allen data (https://portal.brain-map.org/atlases-and-data/rnaseq/mouse-whole-cortex-and-hippocampus-10x) using scrattch.hicat map_sampling function. The parameters were set as: markers.perc = 0.8, iter = 10. To categorize cells among EGFP positive, tdTomato positive, EGFP DE genes positive, and tdTomato DE genes positive cells, we applied tsne on the broad glutamatergic dimensions (**Supplemental Figure 2A**).

### MERFISH tissue preparation and imaging

For fresh frozen tissue preparation, aged 2-3 months mice were anesthetized with 2% isoflurane (v/v); their brains were harvested and embedded with O.C.T (Tissue-Tek), and were then immediately frozen using the combination of 2-Methylbutane (Millipore Sigma) and liquid nitrogen (for details see Tissue Preparation Guide CG000240, 10x genomics). Fresh frozen brains were sectioned into 10-µm-thick slices using a cryostat (MICROM, HM505E). Slices with mPFC region were saved for subsequent hybridization. MERSCOPE Sample Prep Kit (Vizgen) was used to perform *in situ* hybridization on snap frozen brain slice according to the manufacturer’s instruction. Probes for the DE genes of EGFP or tdTomato were selected according to the data of 10x single-cell sequencing. Probes for the layer specific genes were selected based on previous studies^17, 35^. All probes were made by Vizgen (MERSCOPE 140 Gene Panel). The MERSCOPE instrument and the MERSCOPE Analysis Computer (Vizgen) were used to acquire MERFISH raw data. We performed imaging by following the instruction of MERSCOPE Instrument User Guide (Vizgen).

### Pseudo co-localizing analysis of MERFISH data

Because EGFP-labelled MP neurons are rarely found in 10-µm MERFISH slices (∼10/slice), it is not accurate to assess their marker genes by calculating the proportion of MP neurons among marker gene-positive neurons. Therefore, we developed a pseudo co-localizing analysis to determine whether MP neurons and their marker gene-positive neurons are in the same spatial cluster. To extract the spatial variable genes among the DE genes of EGFP neurons, we applied single-cell graph cuts optimization^20^ (scGCO) on mPFC regions of all MERFISH imaged slices. The common spatial variable genes in all repeated slices were selected. The selected gene enriched neurons (gene counts ≥ 1) were marked. Next, we labeled MP neurons on Zeiss imaging slices by using viral injection (see **Surgeries**) and histology (see **Histology for Zeiss imaging**). These Zeiss images were registered to the MERFISH images, which marked by selected gene enriched neurons. The labeled MP neurons in Zeiss images and the selected gene enriched neurons in MERFISH images were further marked by equal sized circles, which acted as pseudo-neurons (Figure 2B). Since the number some selected gene enriched neurons in MERFISH were far more than labeled MP neurons in Zeiss images, we reduced this number by setting the threshold of gene counts (only the neuron with gene counts larger than this threshold can be selected as gene enriched neuron). Threshold settings varied from gene to gene. We set the threshold according to whether the neurons enriched with the selected gene had similar numbers to EGFP neurons. The overlapping pixels between pseudo-neurons from Zeiss and MERFISH images were calculated. DE genes enriched neurons that showed significantly higher overlap than 95 percentile of random genes enriched neurons (**Supplemental Figure 3B**) were chosen for the next step. To further analyze how well the co-localization is, we calculated co-localization performance index, which equals to the number of co-localized pixels/gene count threshold.

To validate this method, we retrieved the mRNA profile of the top three selected genes positive neurons using single-cell sequencing data. Since glutamatergic neurons are dominant (68%) in the MP neurons (**Figure 3C**), we focused only on glutamatergic MP neurons. The ratio of intratelencephalic (IT) / pyramidal tract (PT) neurons and Bcl11b positive IT/PT neurons were calculated. These two ratios were compared among EGFP positive neurons (MP neurons) and selected genes positive neurons. The method of pseudo-co-localizing analysis was validated when the rank of co-localization performance index of selected genes matched the rank of these genes calculated by above two ratios (**Figure 2E**; **Supplemental Figure 3D**).

## Acknowledgement

We thank the general assistance from Sun lab members, including Chunzhao Zhang, Dr. Jiaman Dai, Maycie Schultz, Madeline Bershinsky, Karissa Kiser, Samuel Crouse, Dr. Chase Cao, and Brandon L Robert. We thank Dr. Zhaojie Zhang and Qun Ren for the assistance on scRNA-seq, MERFISH, and microscopy. We thank Dr. Sean M. Harrington for the assistance on R script writing. Dual Lick Port Detector (www.janelia.org/open-science/dual-lick-port-detector) is a gift from Dr. Nuo Li. This work is supported by office of the director, National Institutes of Health (OD), grants from National Institute of Mental Health (1R21MH131363-01), National Institute of Biomedical Imaging and Bioengineering (1R21EB032609-01), National Institute on Aging (R21AG072803), National Institute of General Medical Sciences (2P20GM121310), and National Institute of General Medical Sciences of the National Institutes of Health (P20GM103432).

