## Supplementary Figures for "Control of behavioral uncertainty by divergent frontal circuits"

#### Supplemental Figure 1

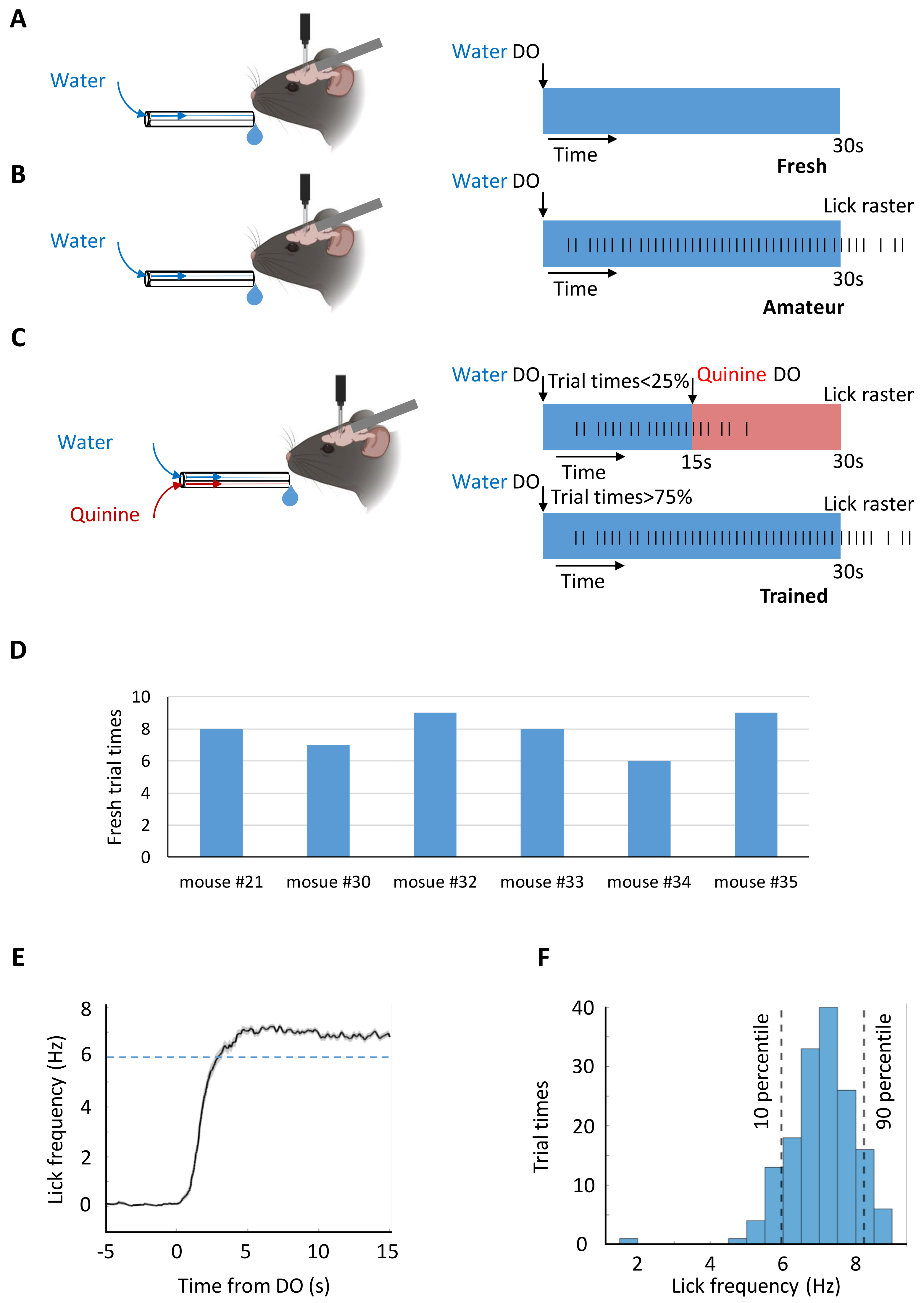

**Supplemental Figure 2**

Procedure of model generation

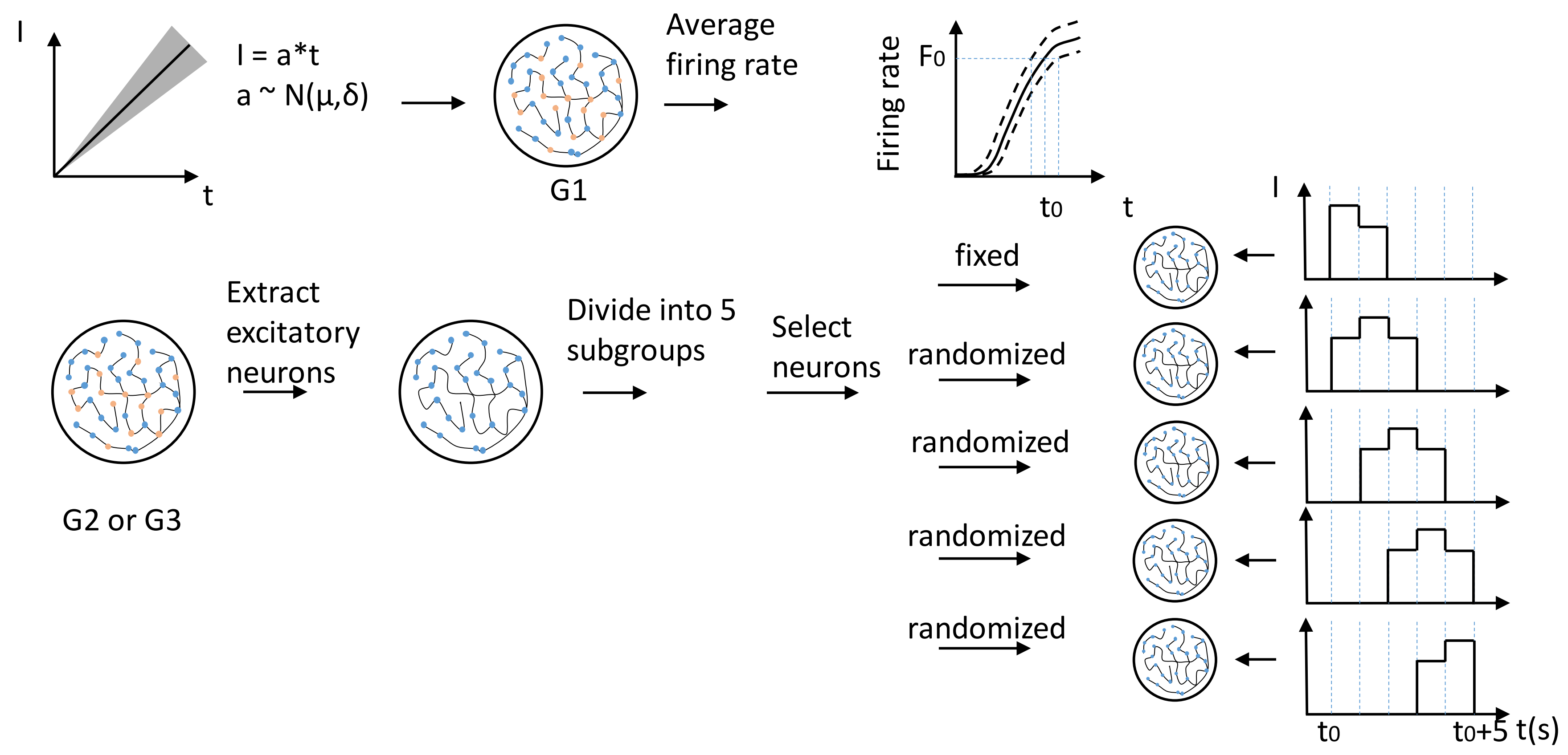

**Supplemental Figure 3**

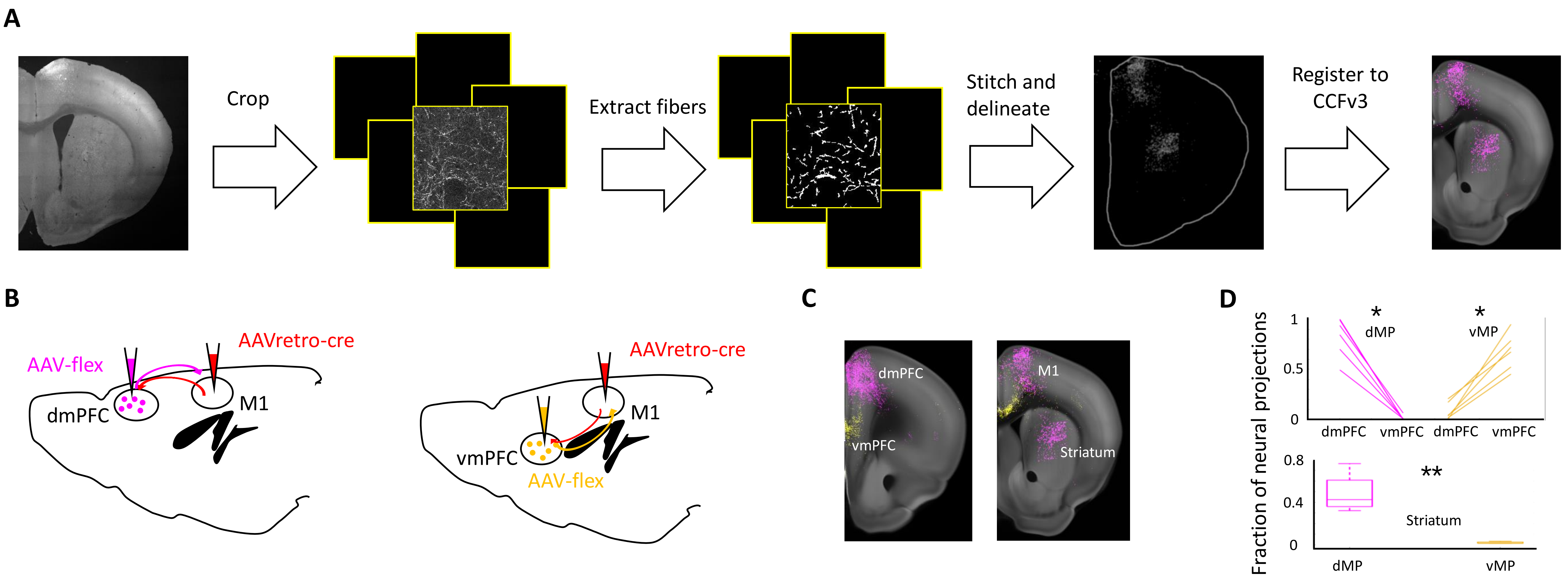

### Supplemental Figure 4

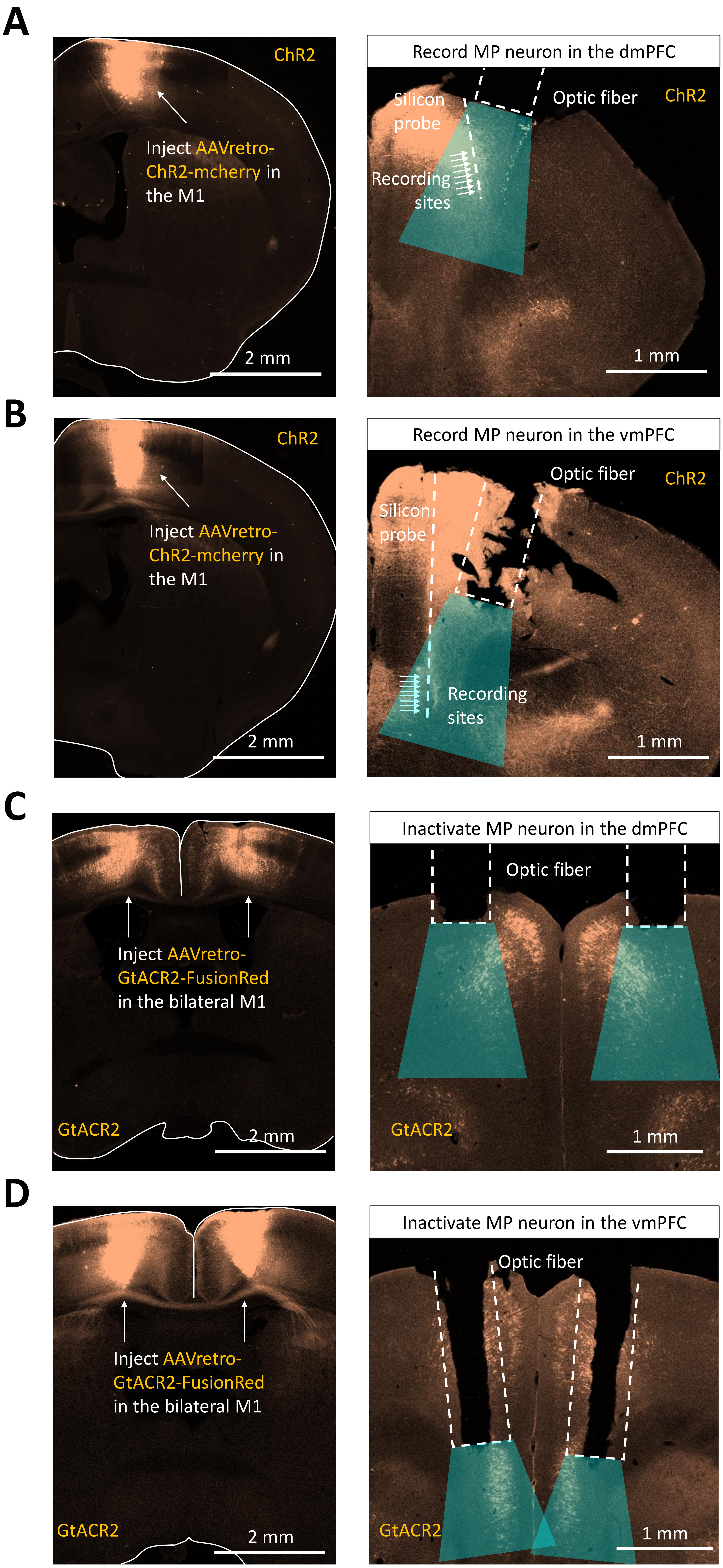

### Supplemental Figure 5

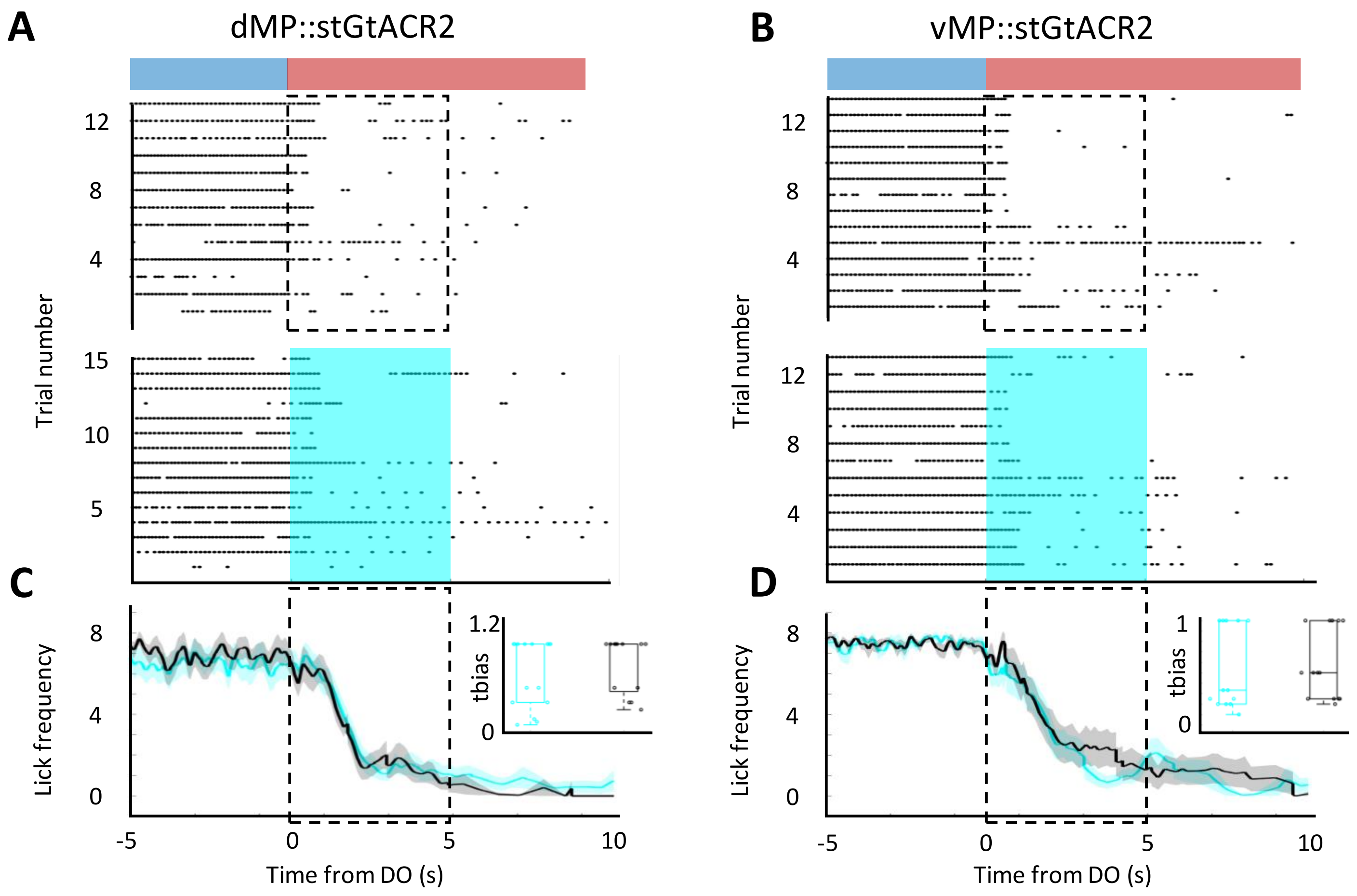

#### Supplemental Figure 6

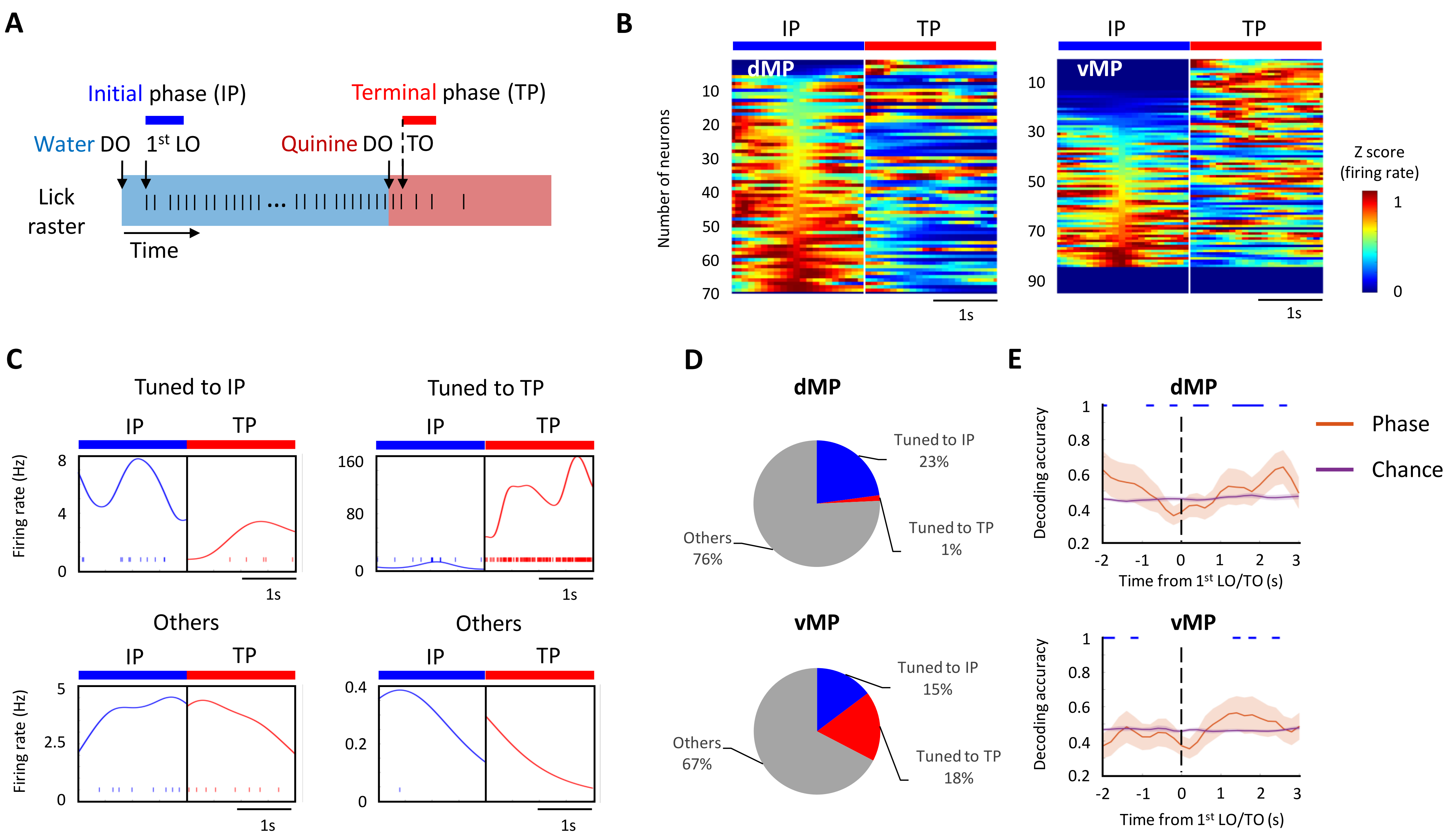

**Supplemental Figure 7**

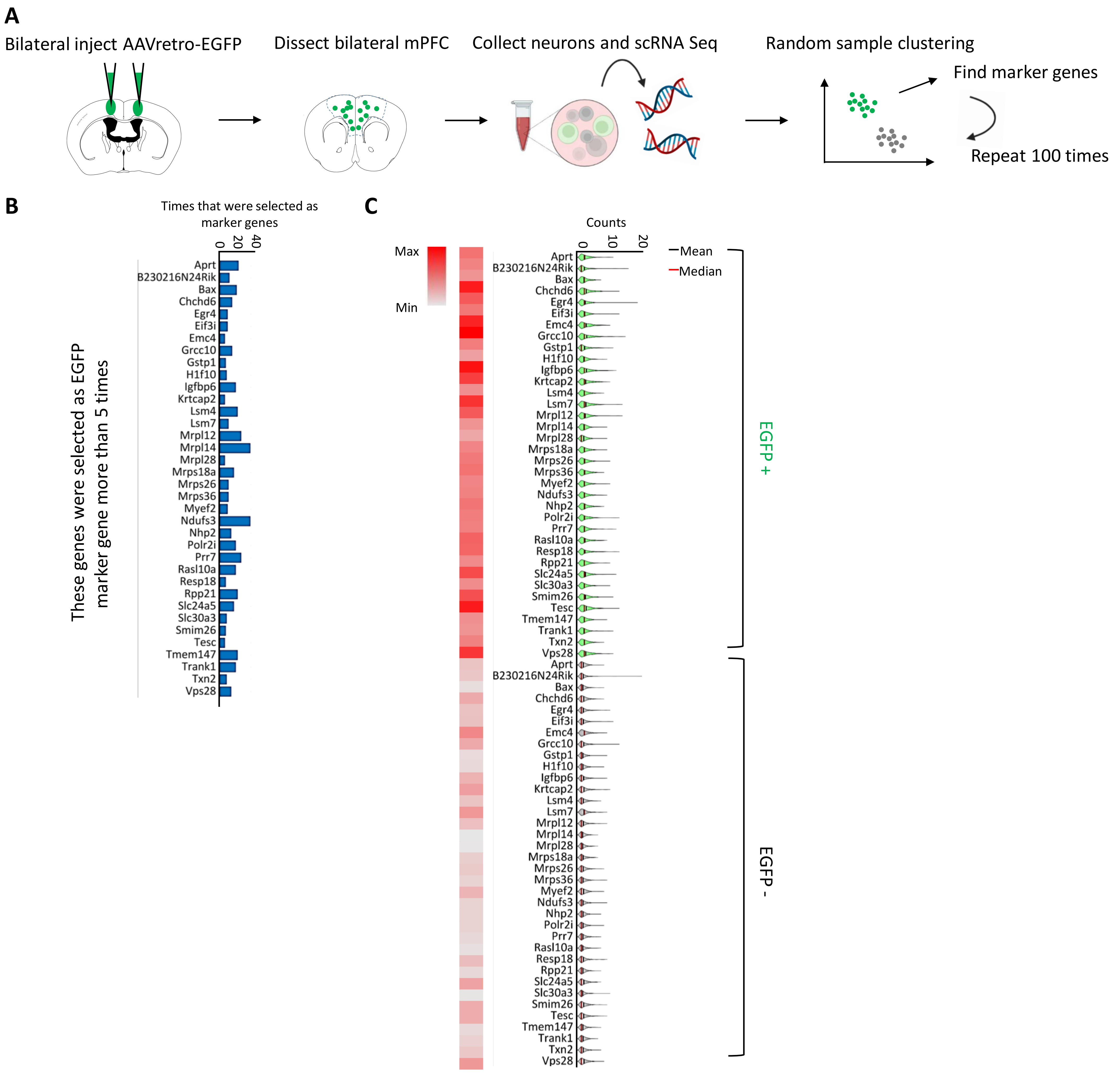

### Supplemental Figure 8

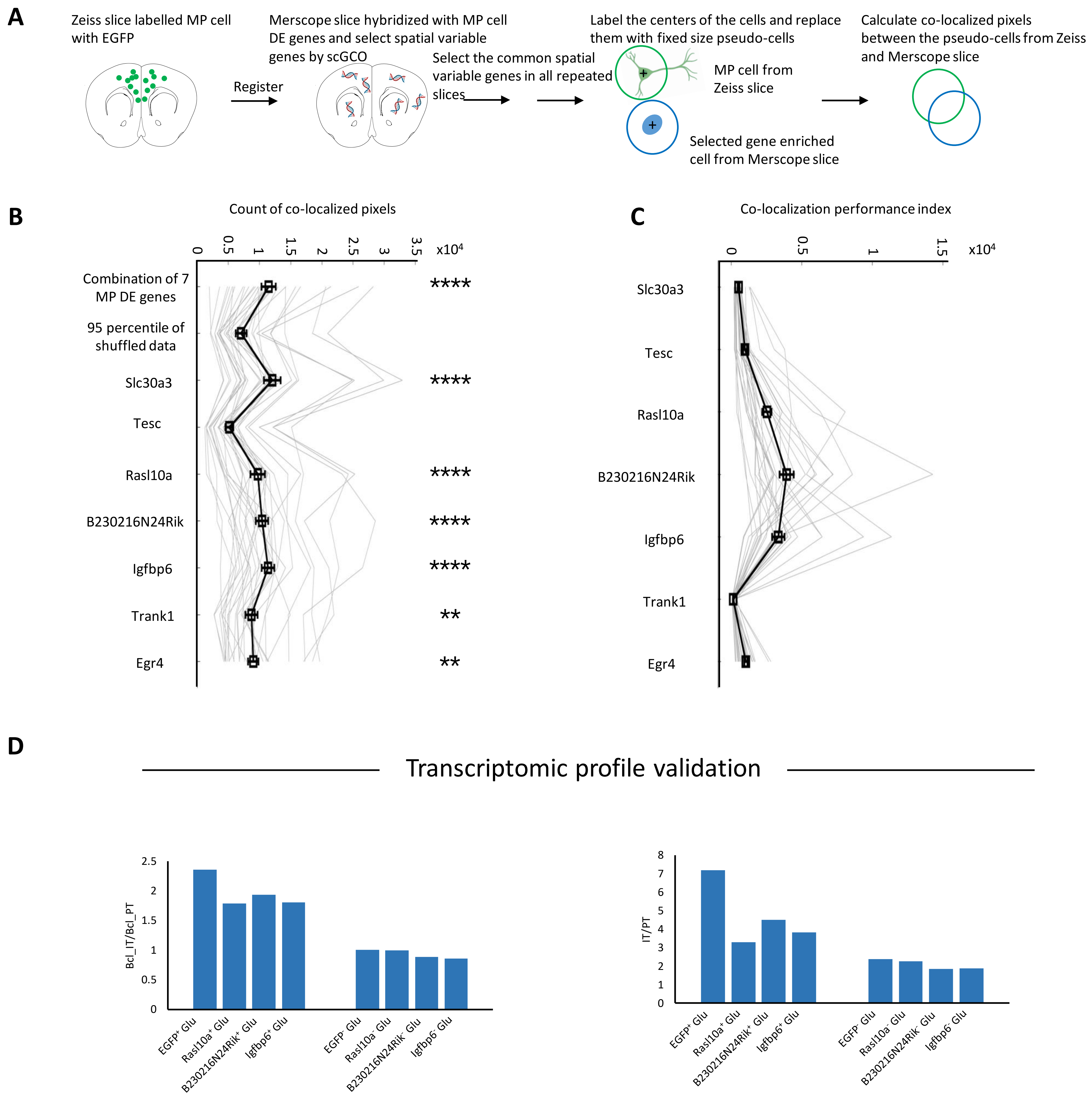

#### Supplemental Figure 9

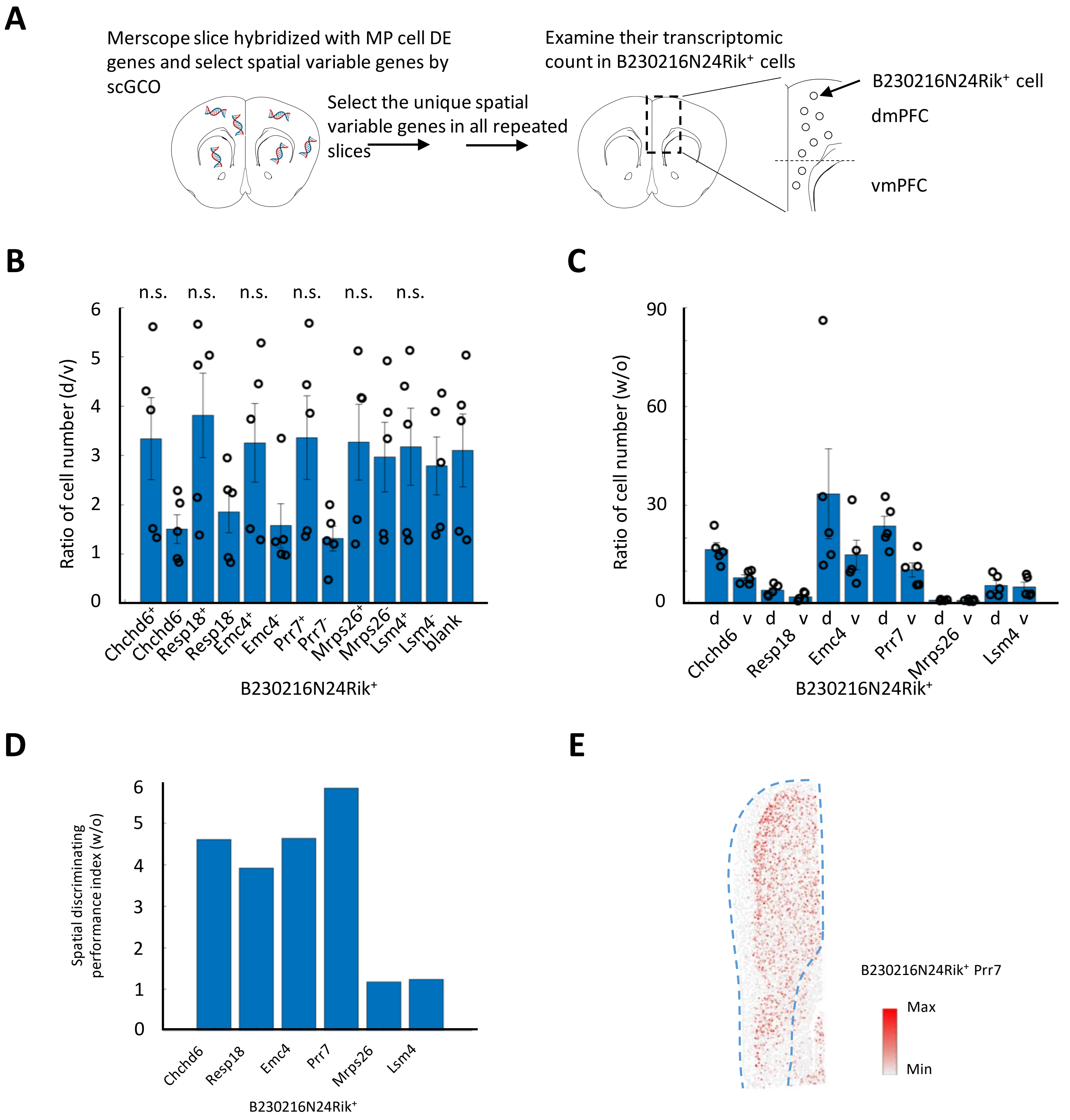

#### Supplemental Figure 10

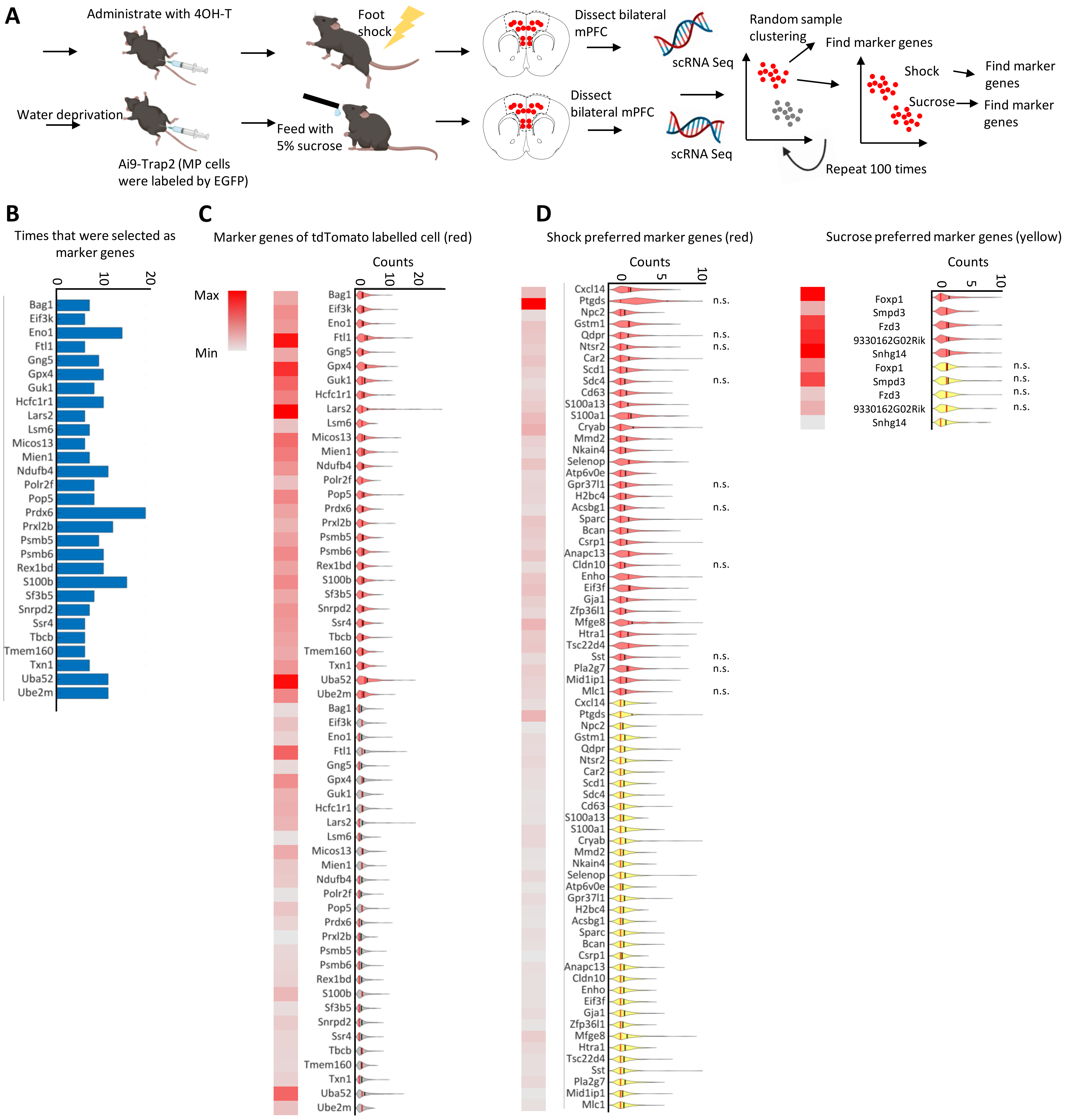

#### Supplemental Figure 11

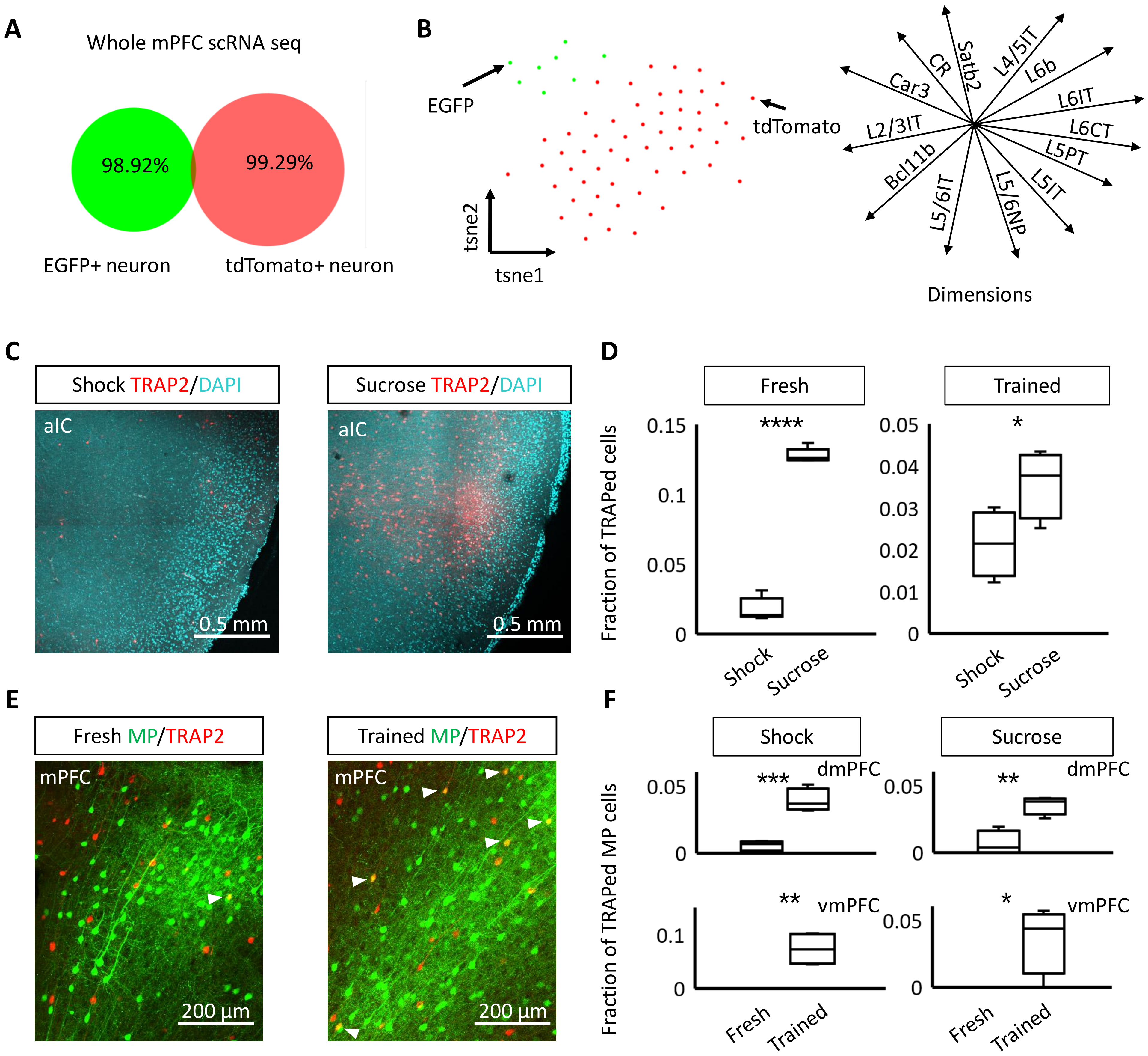

**Supplemental Figure 12**

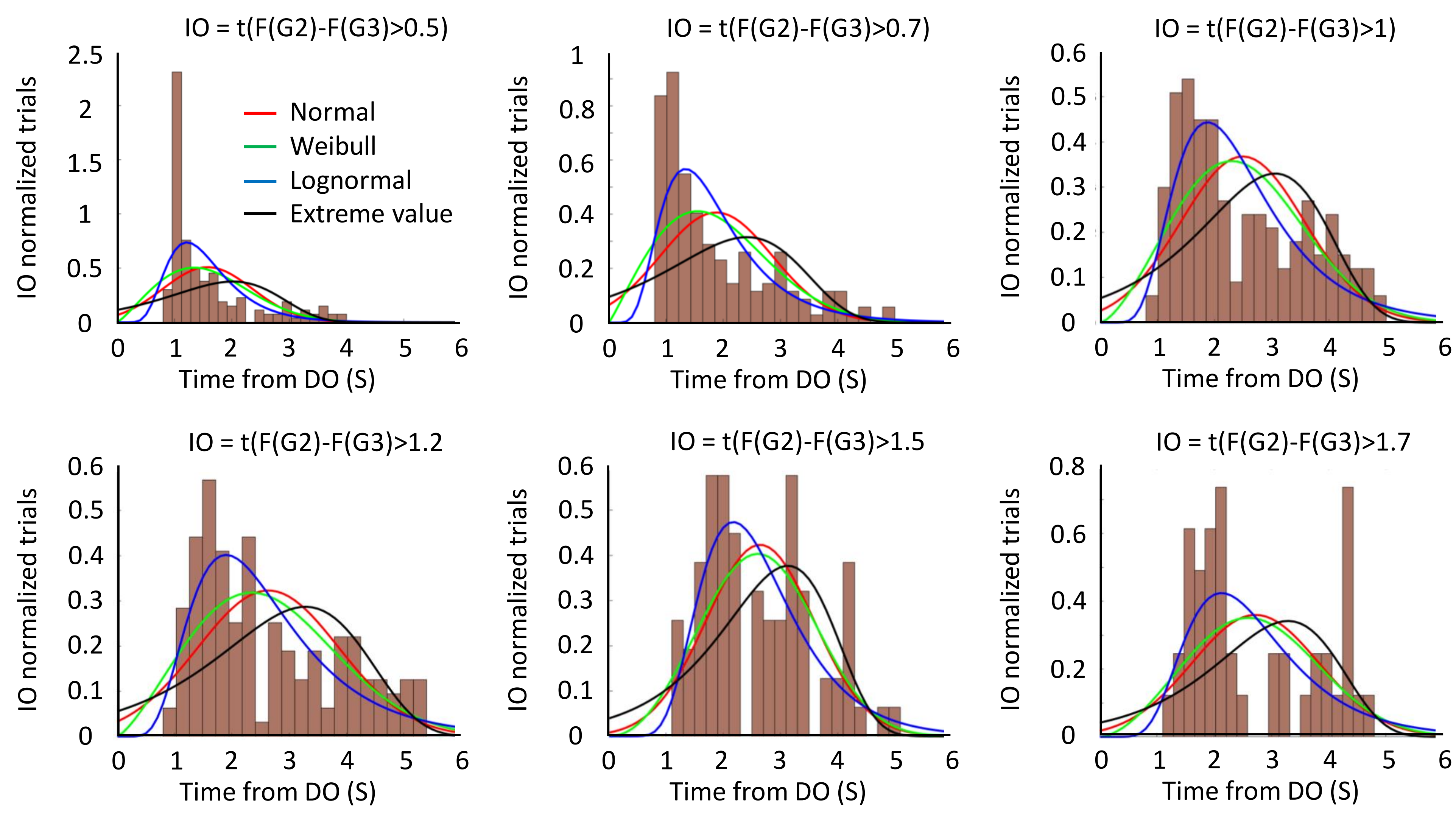
